# Hyperspectral sensing of photosynthesis, stomatal conductance, and transpiration for citrus tree under drought condition

**DOI:** 10.1101/2021.02.26.433135

**Authors:** Jing-Jing Zhou, Ya-Hao Zhang, Ze-Min Han, Xiao-Yang Liu, Yong-Feng Jian, Chun-Gen Hu, Yuan-Yong Dian

**Affiliations:** College of Horticulture & Forestry Sciences, Huazhong Agricultural University, Wuhan 430070, China; Hubei Engineering Technology Research Center for Forestry Information, Wuhan 430070, China; Key Laboratory of Horticultural Plant Biology (Ministry of Education), College of Horticulture and Forestry Science, Huazhong Agricultural University, Wuhan 430070, China

**Keywords:** water stress, photosynthetic CO_2_ assimilation rate, leaf conductance, transpiration rate, hyperspectral reflectance, machine language algorithms

## Abstract

Obtaining variation in water use and photosynthetic capacity is a promising route toward yield increases, but it is still too laborious for large-scale rapid monitoring and prediction. We tested the application of hyperspectral reflectance as a high-throughput phenotyping approach for early identification of water stress and rapid assessment of leaf photosynthetic traits in citrus trees. To this end, photosynthetic CO_2_ assimilation rate (*Pn*), stomatal conductance (*Cond*) and transpiration rate (*Trmmol*) were measured with gas-exchange approaches alongside measurements of leaf hyperspectral reflectance from citrus grown across a gradient of soil drought levels. Water stress caused *Pn, Cond* and *Trmmol* rapid and continuous decreases in whole drought period. Upper layer was more sensitive to drought than middle and lower layers. Original reflectance spectra of three drought treatments were surprisingly of low diversity and could not track drought responses, whereas specific hyperspectral spectral vegetation indices (SVIs) and absorption features or wavelength position variables presented great potential. Performance of four machine learning algorithms were assessed and random forest (RF) algorithm yielded the highest predictive power for predicting photosynthetic parameters. Our results indicated that leaf hyperspectral reflectance was a reliable and stable method for monitoring water stress and yield increasing in large-scale orchards.

**Highlight:** An efficient and stable methods using hyperspectral features for early and pre-visual identification of drought and machine learning techniques for predicting photosynthetic capacity.

## Introduction

Agriculture worldwide accounts for up to 70% of the total consumption of water. Water demand for agriculture used for irrigation will remain the largest and increase by 60% in 2025 (Boretti and Rosa 2019). Global warming is projected to increase evaporation and to reduce soil moisture (Samaniego et al. 2018). Consequently, climate change may exacerbate droughts which may set in more quickly, be more intense and last longer (Trenberth et al. 2014). Fruit trees such as citrus, sensitive to droughts, are already resulting in decreased yields and poor tolerance to pests and stresses (Morgan et al. 2014). Photosynthesis is an important physiological activity in the growth process of green plants and generally limited by soil drought (Zhu et al. 2008). Photosynthetic efficiency is not just connected to potential yield increases but also influences efficiency of the use of resources such as water (Heckmann et al. 2017). Drought often leads to low net photosynthetic rate (Xiao et al. 2019). Improvements in pant photosynthetic efficiency are expected to play a major role in the efforts to increase agriculture productivity (Long et al. 2015, Ort et al. 2015, Osco et al. 2020). An important reason for the insufficient exploration of the potential for changes to water use and photosynthesis for fruit yield forecast and quality improvements is the lack of appropriate high-throughput screening methods.

Further information of fruit trees’ responses to water stress at the large scale throughout the growth period can improve the efficiency of water use. Severe drought has been associated with regional-scale tree mortality and premature senility worldwide which led to reductions in yields (McDowell et al. 2008). Moderate drought stress is one of the commonly method to induce flower buds in tropical and subtropical fruit trees. A moderate water shortage treatment can make trees enter reproductive growth as soon as possible and promote flowering for improving fruit quality and regulating the maturity period (Chica and Albrigo 2013, Li et al. 2017). In addition, plants can be irreversibly affected before visible symptoms of water stress appear(Yordanov et al. 2003). So, a pre-symptomatic or pre-visual detection of plant physiological changes was urgent for avoiding severe damages (Chaerle and Van Der Straeten 2000, Gerhards et al. 2019). Physiological, morphological and biochemical change were observable early at the drought onset (fast changes) or after some time (slow changes) when green plant was encountered soil drought (Acevedo et al. 1971, Zaher-Ara et al. 2016, Sobejano-Paz et al. 2020). Measuring photosynthesis data is a challenge impacted by heterogenetic environmental parameters such as soil moisture content (Zhang et al. 2020). Advancements in phenotyping techniques capable of rapidly assessing the effects of drought on plant photosynthetic responses is necessary to understand plant traits under predicted future environmental conditions (Cotrozzi et al. 2020).The functional responses that are associated with increased yield, such as improving photosynthetic productivity under stressful conditions requires new techniques to quantify this parameter, yet traditional methods rely on leaf sampling and analysis under laboratory conditions or using in-field gas exchange systems (Long and Bernacchi 2003, Osco et al. 2020). This method can provide very precise photosynthetic information but was costly, time-consuming and hard to accomplish especially in citrus-growing mountain.

Remote sensing communities have long used spectrum or spectral vegetation indices to estimate plant biochemical and morphological properties, which also presents huge potential in assessing photosynthetic capacity of plants quickly and non-destructively at different scales (ground, airborne, and satellite) (Asner 1998, Yendrek et al. 2017, Gamon et al. 2019). Hyperspectral spectra ranging from the visible over the near infrared to the intermediate infrared can provide spectral features regarding differences in leaf metabolism, structure, physiological and chemical traits in associated plant conditions (Pinter et al. 2003, Cotrozzi et al. 2017, Santoso et al. 2019, Osco et al. 2020, Streher et al. 2020). It is very popular to use hyperspectral reflectance to assess crop physiological and biochemical parameters. Nutritional status (Bruning et al. 2019, Chen et al. 2019), chlorophyll or carotenoid contents (Sonobe et al. 2020a, Sonobe et al. 2020b, Yamashita et al. 2020), water content (Kong et al. 2019, Chen et al. 2020), heavy metal content (Zhou et al. 2020c) and species composition (Manevski et al. 2011) in crops have been estimated by using leaf reflectance spectrum. Application of hyperspectral spectra to assess plant function or physiology is complex, as the mechanisms linking spectra reflectance and emission to plant functional traits are not always clear or known (Gerhards et al. 2019). Spectroscopy techniques coupled with deep learning algorithms has been also used for leaf morphological and biochemical traits with the highest photosynthetic potential (Serbin et al. 2014, Osco et al. 2020). Photosynthetic capacity of crop plants has been evaluated based on leaf reflectance successfully by using specific wavelengths or indices related to photosynthesis status of the plants over a wide range of species (Driever et al. 2014, Garriga et al. 2017, Heckmann et al. 2017, Silva-Perez et al. 2018, Fu et al. 2019, Meacham-Hensold et al. 2019, Fu et al. 2020, Watt et al. 2020, Zhang et al. 2020). Although, further research on citrus is necessary. Macro- and micronutrient content of citrus had been successfully predicted with leaf reflectance data (Osco et al. 2020). Very little studies have focused on using leaf reflectance spectra to monitor the respond of citrus leaves to water stress and estimate the photosynthetic capacity.

Citrus is the most widely cultivated fruit crop worldwide and also abundant within China. In the past 20 years, citrus industry has developed rapidly in the world. The most outstanding research about citrus was related to molecular breeding, stress respond and post-harvest treatment (Wu et al. 2016, Ling et al. 2017, Liu et al. 2017, Wang et al. 2017). In this study, citrus leaves—more specifically, from lemon (*Citrus limon*) trees—were selected to compose the experimental dataset. Measurements of leaf reflectance and photosynthesis were taken from an experiment that included a factorial water stress applied to citrus trees. Using this data, we address the following question: (1) how citrus physiology (leaf *Pn, Cond* and *Trommol*) respond to continuous soil water stress and re-watering (2) what is the variation in the leaf reflectance spectra of citrus in different drought treatment? what is the most stable hyperspectral information selected responding citrus leaves to water stress? (3) how leaf hyperspectral reflectance spectra can best be used to predict photosynthetic capacity of citrus leaves? The answers to these questions will help to facilitate the spectral responds of citrus to drought and photosynthesis prediction models.

## Materials and Methods

### Experimental design

The study was conducted at the greenhouse facility of Huazhong Agriculture university using lemon (*Citrus limon*) as the selected plant material, which is located in Wuhan, Hubei province, China (113°41′-115°05′E, 29°58′-31°22′N). Wuhan is one of the largest cities on the upper and middle reaches of the Yangtze River in central China. The annual average temperature, mean annual relative humidity, precipitation and annual average frost-free period are 16.9 °C, 77%, 240 days, 1259 mm, respectively (Yan et al. 2019). Random block design was used in this study. 4-year-old lemon trees (Femminello) propagated by bud grafting to trifoliate orange rootstocks were used. These trees ranged in height from 2 to 2.5 m growing in 60-cm plastic pots containing potting mix of commercial medium and perlite (3:1). Trees in the greenhouse were exposed to natural variations in photoperiod throughout the experiment during Summer (from August to September) 2020. Each level included 8 lemon trees. The soil moisture was controlled to 35% (Normal water supply), 25% (Mild stress), 15% (Moderate stress) and 10% (Severe stress). All the trees were firstly adopted to drought treatment for 21days in 6-Aug, 12-Aug, 18-Aug, 26-Aug and then watered three times in 26-Aug, 4-Sep and 9-Sep. Three trees were randomly selected from each drought treatment, and two randomly selected leaves from the upper, middle and lower layers of each selected trees were used for measuring photosynthetic and spectral related parameters in 6-Aug (day 1), 12-Aug (day 7), 18-Aug (day 13), 26-Aug (day 20), 4-Sep (day 28) and 9-Sep (day 33).

### Hyperspectral Measurement Processing

The spectral radiance of the lemon leaves was measured with a full-range hyperspectral PSR-3500 spectroradiometers. The FieldSpec collects data in the 350–2500 nm spectral range, with a resampled spectral resolution of 1nm before 1006nm and 3.5 nm after 1006nm. Leaf reflectance data was collected on the surface of the leaf at 2 positions per leaf using the leaf clip from mature leaves. Ten measurements were conducted in each leaf position to produce one mean spectral reflectance. Before each spectral measurement, a white surface plate was registered to calibrate the equipment and convert the digital number to a physical signal (Osco et al. 2020). Leaf reflectance was computed as the ratio of leaf radiances relative to the radiance from the white reference panel (Shah et al. 2019).

### Photosynthetic Measurement

Immediately after the spectral reflectance scan, the selected leaf was placed into the leaf room of the LICOR 6400XT gas analyzer (LICOR Biosciences, Lincoln, NE, United States). Measurements were initiated at a saturating light (1000 mmol m^−2^ s^−1^), a block temperature of 25L, and a flow rate of 500 mmol mmols^-1^. leaf photosynthetic CO_2_ assimilation rate (*Pn*, μmol CO_2_ m^-2^s^-1^), leaf stomatal conductance (*Cond*, mol HO_2_ m^-2^s^-1^), leaf transpiration rate (*Trmmol*, mmol HO_2_ m^-2^s^-1^) for each leaf was captured after an adjustment period of approximately 30 minutes.

### Extraction of Vegetation Indices

A database of 20 narrow-band spectral vegetation indices (SVIs) (Table 1), which have shown potential for assessing attributes of vegetation parameters related to plant physiology, morphology and biochemistry, were preselected for analysis. They simplified the interpretation of complex vegetation reflection characteristics based on the indirect relationship between plant physiological and structural parameters (Gerhards et al. 2019). Normalized difference vegetation index (NDVI), Ratio vegetation index (RVI), Enhanced Vegetation Index (EVI) can represent greenness, vegetation cover, biomass, LAI and fraction of photosynthetic active radiation (Sobejano-Paz et al. 2020). Greenness Index (GI), Red Edge model (CI_730_), Red Edge model (CI_709_), Chrollophy Index at Green band (CIG), Normalized Difference Red Edge (NDRE), Red and Green Vegetation Index (RGVI), Chlorophyll Absorption Ratio Index (CARI), MERIS Terrestrial Chlorophyll Index (MTC) were related to plant pigments. Because of the short-term changes of xanthophyll under stress, photochemical reflectance index (PRI), and its derivations (i.e. photochemical reflectance index improved, PRI2) was directly related to photosynthesis (Ballester et al. 2018). Moisture Stress Index (MSI), Water Index (WI), Normalized Multi-band Drought Index (NMDI), Global Vegetation Moisture Index (GVMI), Normalized Difference Water Index (NDWI1200), NDWI1240, NDWI1640 were calculated as proxy of water content. All of the data processing and calculations of the SVIs were performed in the python 3.7 software package.

**Table 1:**
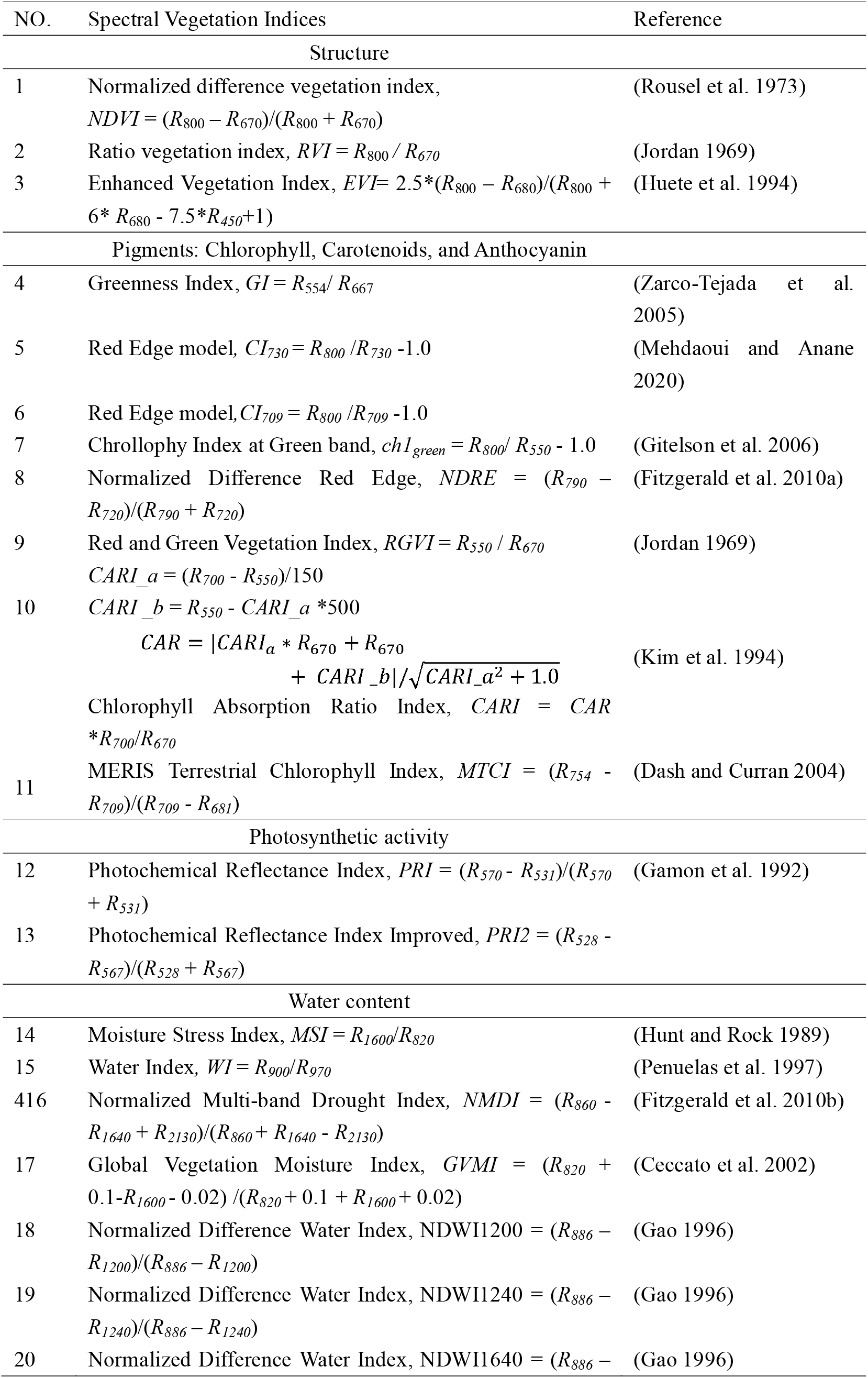
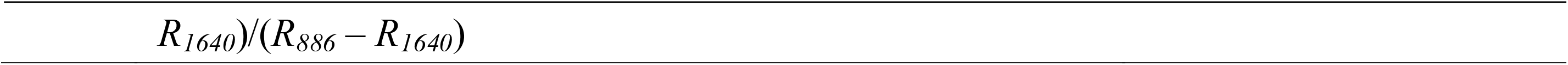
The 20 selected spectral vegetation indices examined in this research, together with their band-specific formulations, and associated principal reference.

We also conducted 40 spectral absorption features and wavelength position variables acquired from leaf spectral reflectance to analysis the different stress level on lemon leaves. The red edge optical parameters from a plant spectrum between 670 nm and 780nm were commonly used in analysis plant. In this paper, we used red edge position (REP) parameter, which is a wavelength position variable indicating biophysical and biochemical parameters of vegetation. The inverted-Gaussian (IG) model was used to extract the red edge optical parameters. The spectral shape of the red edge reflectance can be modeled as equation (1):

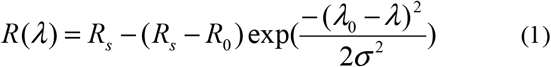

Where *R_s_* is the maximum spectral reflectance; *R*_0_ and λ_0_ are the minimum spectral reflectance and corresponding wavelength, respectively; λ is the wavelength; and σ is the Gaussian function variance parameter. The REP is calculated as equation (2):

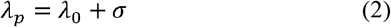

We also acquired the absorption features in 1230 nm to 1650 nm and 1800 nm to 2200nm, as these two spectral ranges are highly corresponding the water stress. The absorption features in a reflectance spectrum include wavelength position (nm), depth, width, area, asymmetry and spectrum absorption index with a continuum removal procedure (Pu et al. 2003). Figure 1 showed a part of typical spectrum (1200 nm - 1300 nm) of lemon leaf to illustrate the feature parameters. The absorption position (P) marks the wavelength at the deepest absorption. The width (W) defines the full width half maximum. X_1_ and X_2_ is the wavelength of left and right shoulder at the position of the full width half maximum. Δλ which represent the value of W is calculated by X_2_-X_1_. Y is the corresponding reflectance of X_1_ and X_2_. The absorption depth (DEP) is the depth of the feature minimum relative to the hull. The absorption area (Area) is the area of the absorption district. The asymmetry of an absorption feature derived as the ratio of the left area (label A in Figure 1) of the absorption center to the right area of the absorption center. L is tangent line and the slope of L can be calculated. Spectrum absorption index (SAI) defines the absorption intensity, which was calculated as equation (3):

**Figure 1.**
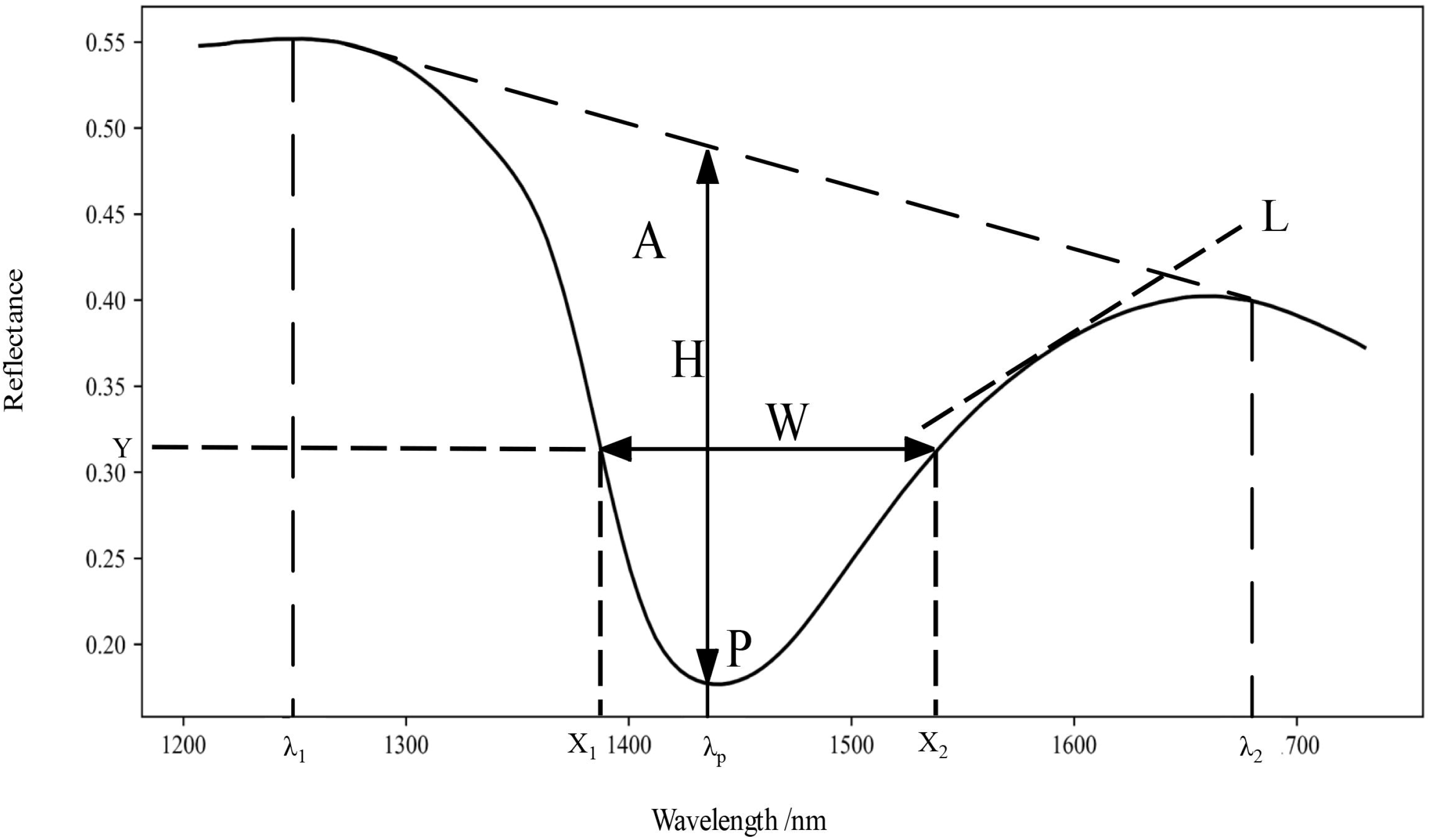
A part of lemon leaf reflectance (1200-1700nm) and definitions of absorption features (where H means absorption depth, W represents the full width half maximum, P is the wavelength of absorption position, A is the area left area of the absorption center, L is tangent line, X_1_ and X_2_ is the wavelength of left and right shoulder at the position of the full width half maximum. Y is the corresponding reflectance of X_1_ and X_2_.)

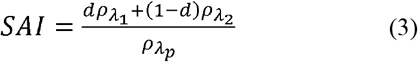

Where λ_l_ and λ_2_ are shoulder wavelength, ρ λ_1_ and ρ λ_2_ are the reflectance at corresponding wavelengths, respectively. *d* is the normalized weight, calculated as equation (4):

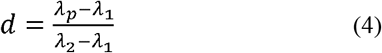

### Principal component analysis (PCA) and ANOVA analysis

In order to understand the difference of lemon leaf reflectance spectra of four drought treatments, PCA was conducted in python 3.7 with the sklearn package. A one-way ANOVA was performed to assess the effect of drought treatment to spectral parameters. Prior to ANOVA analysis, all the data were tested for normality. Differences in means were regarded to be significant if the *P* value was less than 0.05.

### Machine Learning Algorithms

Radom Forest (RF), Support Vector Machines (SVM), Gradient boost (GDboost) and Adaptive Boosting (Adaboost) methods were applied to estimate the *Pn, Cond* and *Trmmol* value. The field data was randomly divided into training (70%) and testing (30%) data. The prediction metrics to evaluate above-mentioned algorithms were the coefficient of determination (R^2^), root-mean-squared error (RMSE) and mean absolute error (MAE). To determine the relationship between the predicted and measured values, the overall model will be evaluated in the regression graph. All the algorithms were implemented in the scikit-learn 0.22.2 package in python 3.7 (Pedregosa et al. 2011).

*Ntree* (i.e., to the number of variables) and *Mtry* (i.e., to the number of variables to randomly sample as candidates at each split) are two key parameters influencing robustness of RF algorithms (Zhang et al. 2018, Zhou et al. 2020b). These two parameters were often set default values (Wang et al. 2016, Zhao et al. 2019).

SVM use a nonlinear kernel function to project input data onto a high dimensional feature space, where complex non-linear patterns can be simply represented (Mountrakis et al. 2011). The key to SVM is the kernel function. Low-dimensional space vector sets are usually difficult to divide. The best choice is to map them to high-dimensional spaces. The classification function of the high-dimensional space can be obtained by selecting the appropriate kernel function (Were et al. 2015). Gaussian radial basis kernel function of the form was applied in this study (Rodriguez-Galiano et al. 2015).

Boosting method establishes several basic estimators (decision tree is used in this paper), each of which can learn to correct the prediction error of prior model in the model sequence, such as GDboost and Adaboost method. The GDboost Algorithm tries to match the residual error of the new predictor with the previous one, while the Adaboost algorithm corrects the unfitness of the training instance through the previous training. The main difference between GDboost and Adaboost was how they deal with the underfitted values (Zhou et al. 2020a). The number of trees (*ntree*) and learning rate (*learning_rate*) were tuned in AdaBoost and GDBoost models.

## Results

### Photosynthetic response to water stress

The sensitivity of *Pn, Cond* and *Trmmol* to water stress of upper layer, middle layer, and lower layer were compared in drought treatment period (i.e. day 1, day 7, day 13 and day 20) and re-watering period (i.e. day 28 and day 33) (Figure 2, supplementary table S1-S2). Comparing among soil water status from 35% (normal water supply) to 10% (severe drought) showed decreases in average *Pn, Cond* and *Trmmol* in upper, middle-, and lower-layers drought treatment period. After re-watering, *Pn, Cond* and *Trmmol* values of different water stress were nearly not significantly different. *Pn, Cond and Trmmol* of normal water supply of upper layer was significantly higher than other water stress in the whole drought treatment time, whereas these values of severe drought was significantly lower than other water stress. Photosynthetic changes with drought in the middle and lower layer were less sensitive than upper layer. *Pn, Cond* and *Trmmol* of moderate or severe drought were lower significantly than normal water supply, whereas these photosynthetic parameters were nearly not significant different of mild drought comparing to normal water supply.

**Figure 2.**
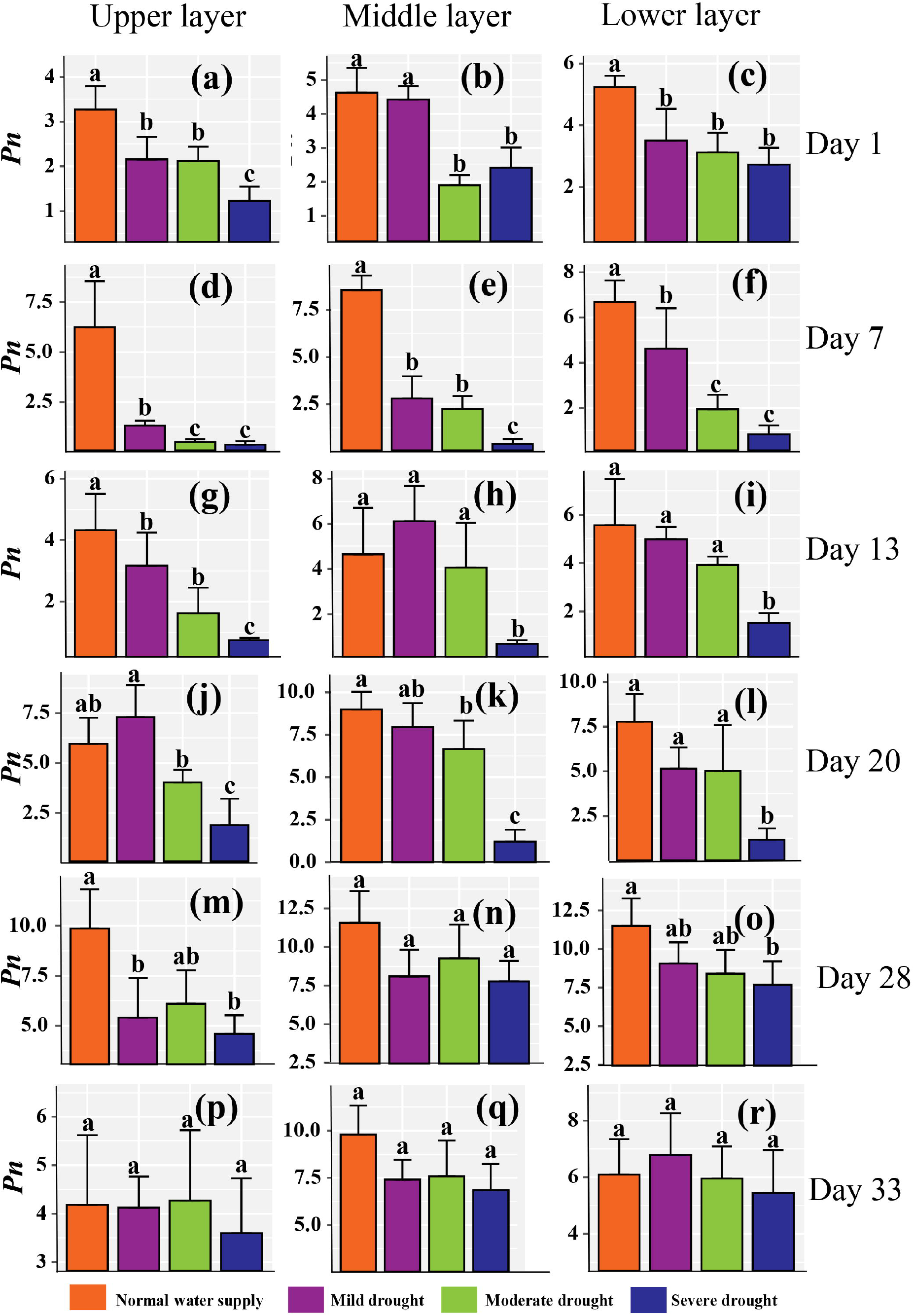
One-way ANOVA test results of photosynthetic CO_2_ assimilation rate (*Pn*, mol m^-2^s^-1^) of upper layer, middle layer, and lower layer of different water stress in drought treatment period (i.e. 6-Aug, 12-Aug, 18-Aug, and 26-Aug) and re-watering period (i.e. 4-Sep and 9-Aug). The data was presented in the form of mean ± standard error, significant differences were indicated by different letters in the same subfigure. Sub-figure (a), (b), (c), (d), (e), (f), (g), (h), (i), (j), (k), (l), (m), (n), (o), (p), (q), (r) were upper layer-6-Aug, middle layer-6-Aug, lower layer-6-Aug, upper layer-12-Aug, middle layer-12-Aug, lower layer-12-Aug, upper layer-18-Aug, middle layer-18-Aug, lower layer-18-Aug, upper layer-26-Aug, middle layer-26-Aug, lower layer-26-Aug, upper layer-4-Aug, middle layer-4-Aug and lower layer-4-Aug.

### Variation of leaf reflectance spectra of different water stress and tracking of leaf hyperspectral reflectance to water stress

Spectra of four drought treatment was of low diversity, percentage of 97.55% of variance was contained in the first three principal components (Figure 3). The reason for this becomes apparent in the correlation matrix of the spectra (Figure 4). Five main independent wavelength ranges have been identified, within which the measurements are closely related. Two correlated ranges are found in the visible spectrum (from about 400 to 480 nm and from 500 nm to 660 nm) and three in the infrared region (from about 720 nm to 1400 nm, 1450nm to 1800nm and from 1900 nm to 2500 nm). Three of them could be reflected in the three main components (Figure 4).

**Figure 3.**
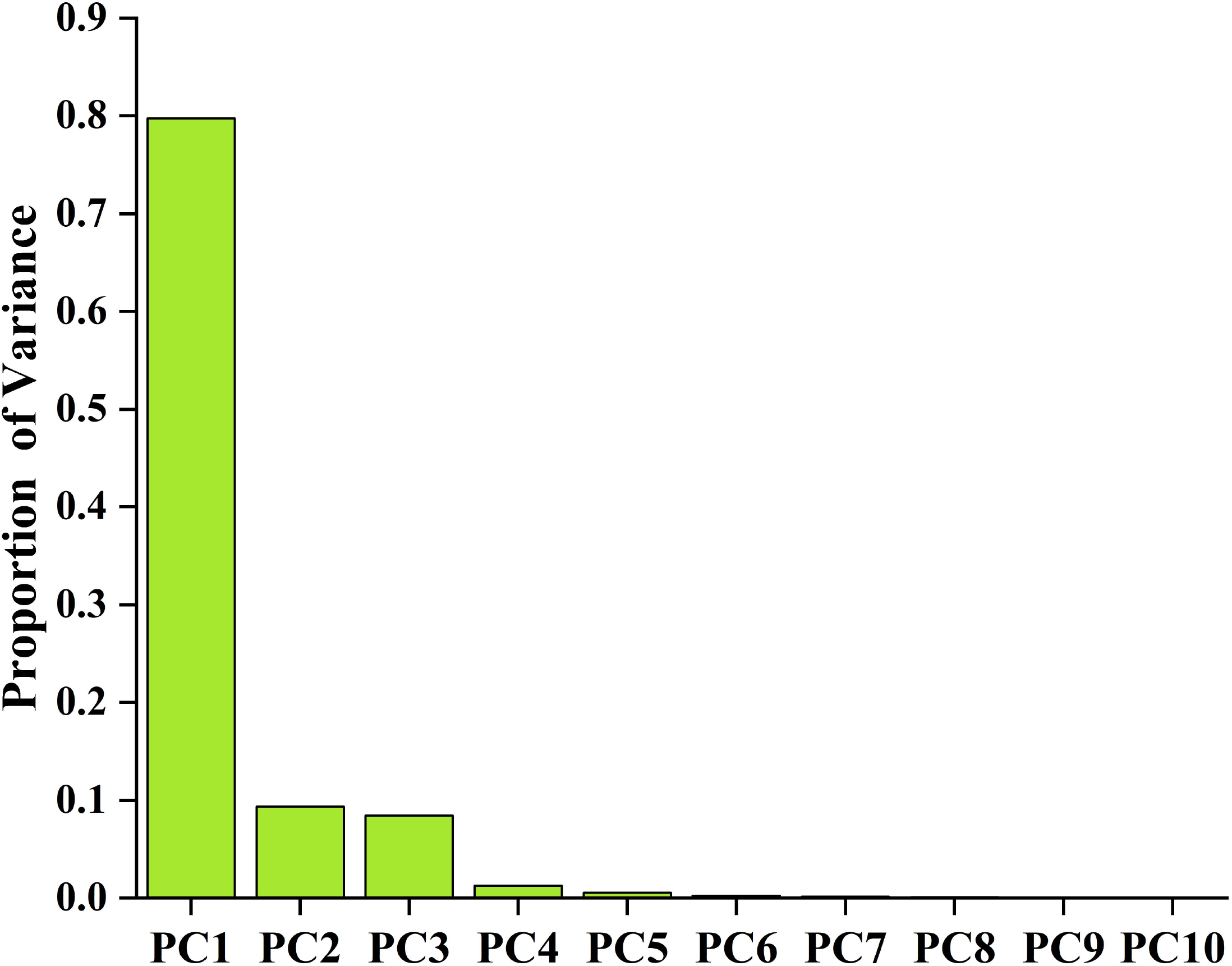
Proportion of variance covered by the first 10 principal components of reflectance spectra in lemon of different drought treatments.

**Figure 4.**
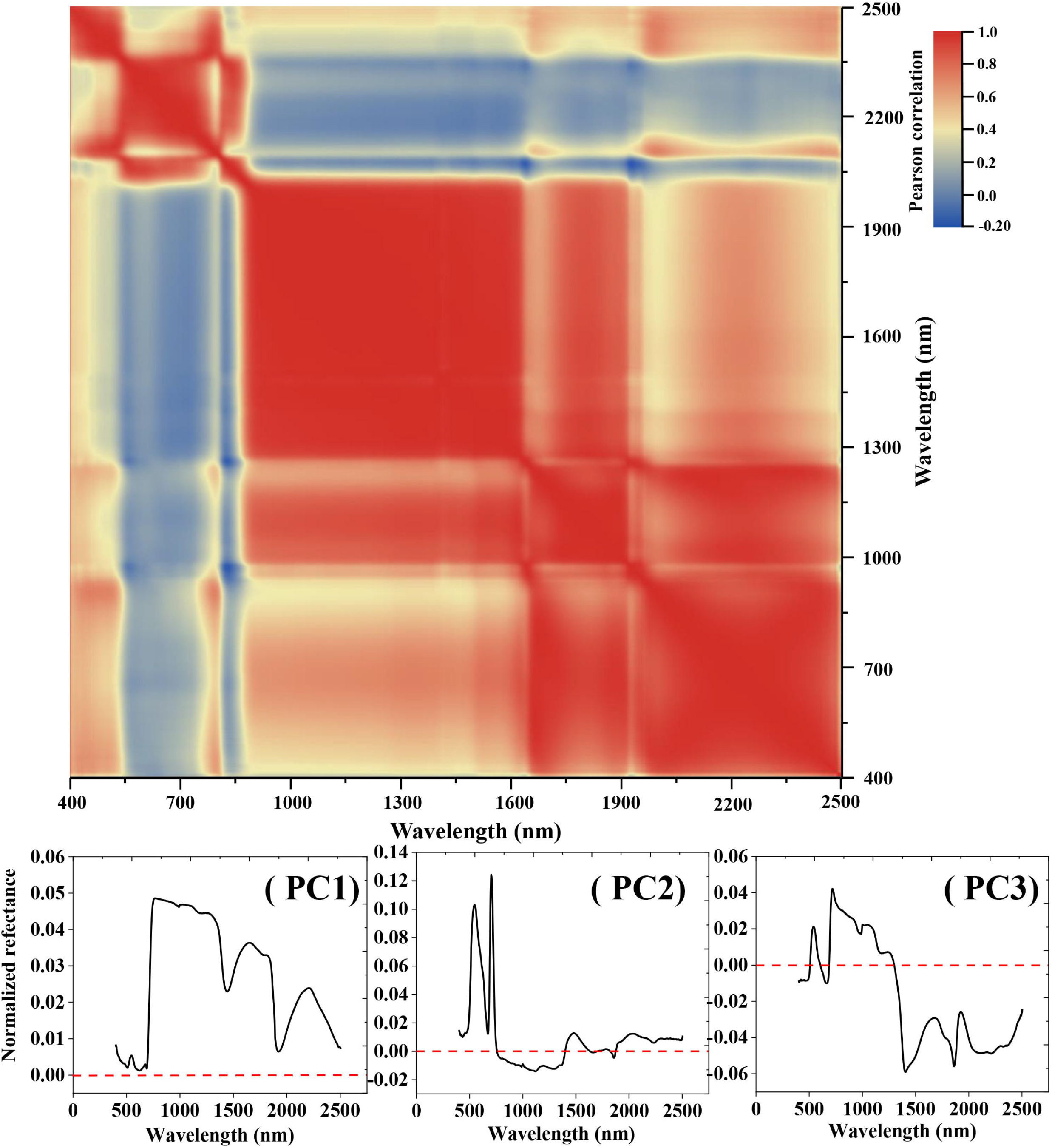
Heat maps of the reflectance spectra of lemon leaves with four drought treatments. Each point shows the Pearson correlation of reflectance at both wavelengths. The panel below shows the three components that account for the largest proportion of variance.

Eighteen hyperspectral parameters selected from sixty parameters presented significant difference among different water stress. PRI, NDVI, RVI, GI, C, NMDI, VIS-λ _*p*_, SW1-fwhmX_1_ and SW1-fwhmX_2_ showed difference just after drought treatment. Especially, PRI was more sensitive to water stress than other hyperspectral parameters and the values of different drought treatments have significant differences in the whole drought period. PRI value of normal water supply was significantly higher than mild, moderate and severe treatment (Figure 5a). NDVI and GI of normal water supply and mild treatment were significantly higher than moderate and severe treatments in day 1 and day 7 (Figure 5b and Figure 5d). RVI shows the opposite trend (Figure 5c). In day 13 and day 20, NDVI and RVI of severe treatment were significantly higher or lower than other treatments (Figure 5b and Figure 5c). GI and C could distinguish the normal water supply, mild or moderate and severe treatment effectively, however, they could not reveal the spectra difference between mild and moderate treatments (Figure 5d and Figure 5e). NMDI, VIS-λ, SW1-fwhmX1, SW1-fwhmX_2_ were effectively used to distinguish severe drought treatment based on spectra from day 1 to day 20 (Figure 5f-5i).

**Figure 5.**
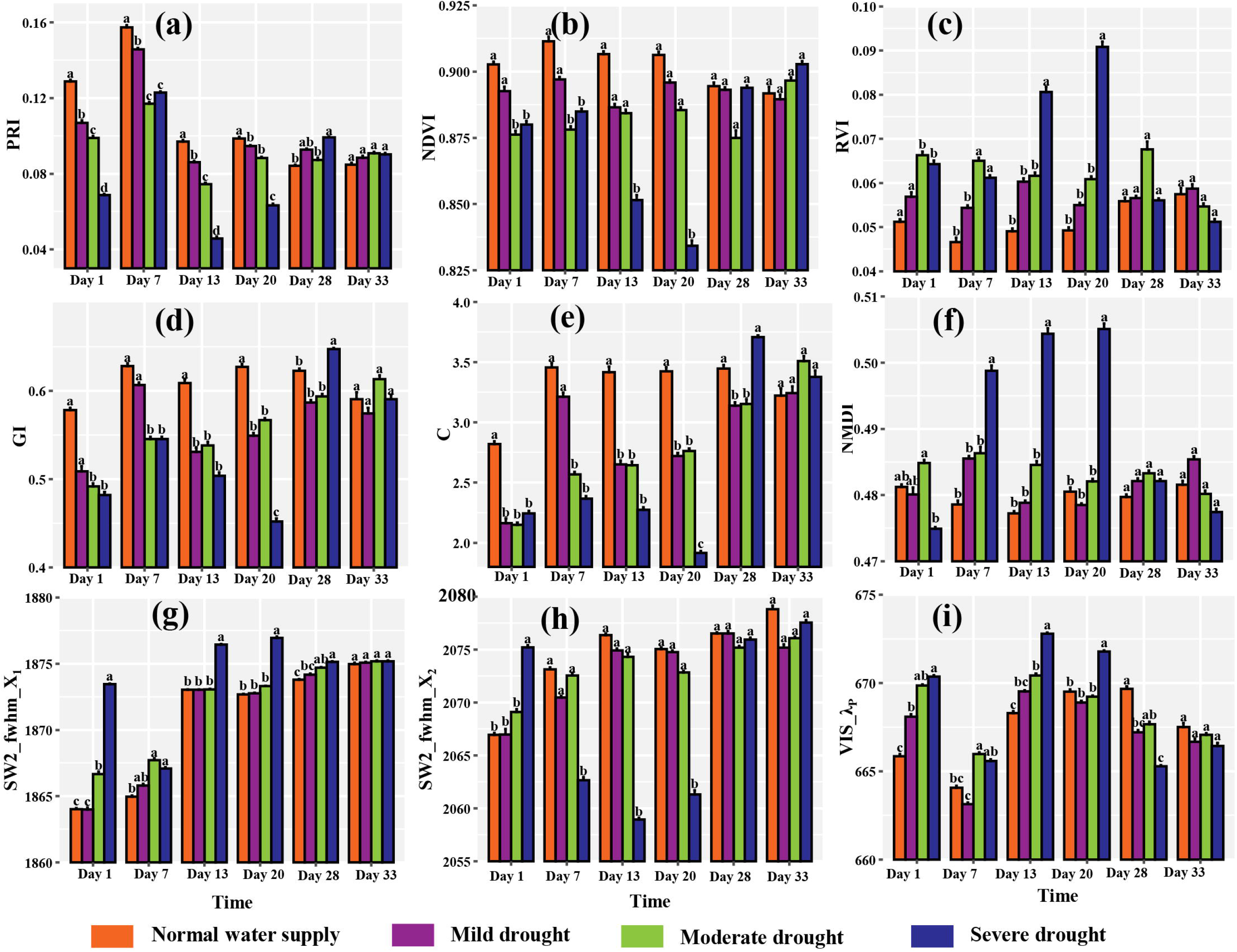
Performance of spectral parameters for revealing the difference of four drought treatments. a, b, c, d, e, f, g, h and i were indicated as PRI, NDVI, RVI, GI, C, NMDI, SW2-fwhm-X_1_, SW2-fwhm-X_2_ and VIS_λ_P_ respectively.

SW1-fwhmY, SW1-fwhm-Δλ, SW1-SAI, SW2-fwhm-Y, SW2-fwhm-Δλ, SW2-SAI, SW2_Area, MSI, NDWI1640, GVMI were useful to distinguish severe drought treatment in day 7, day 13 and day 20. These values of normal water supply, mild and moderate treatment were not significantly different (Figure 6). SW1-fwhmY, SW2-fwhm-Y, MSI values of severe drought were significantly higher than other three treatments, while SW1-fwhm-Δλ, SW1-SAI, SW2-fwhm-Δλ, NDWI1640, SW2_Area, SW2-SAI and GVMI presented opposite trends (Figure 6).

**Figure 6.**
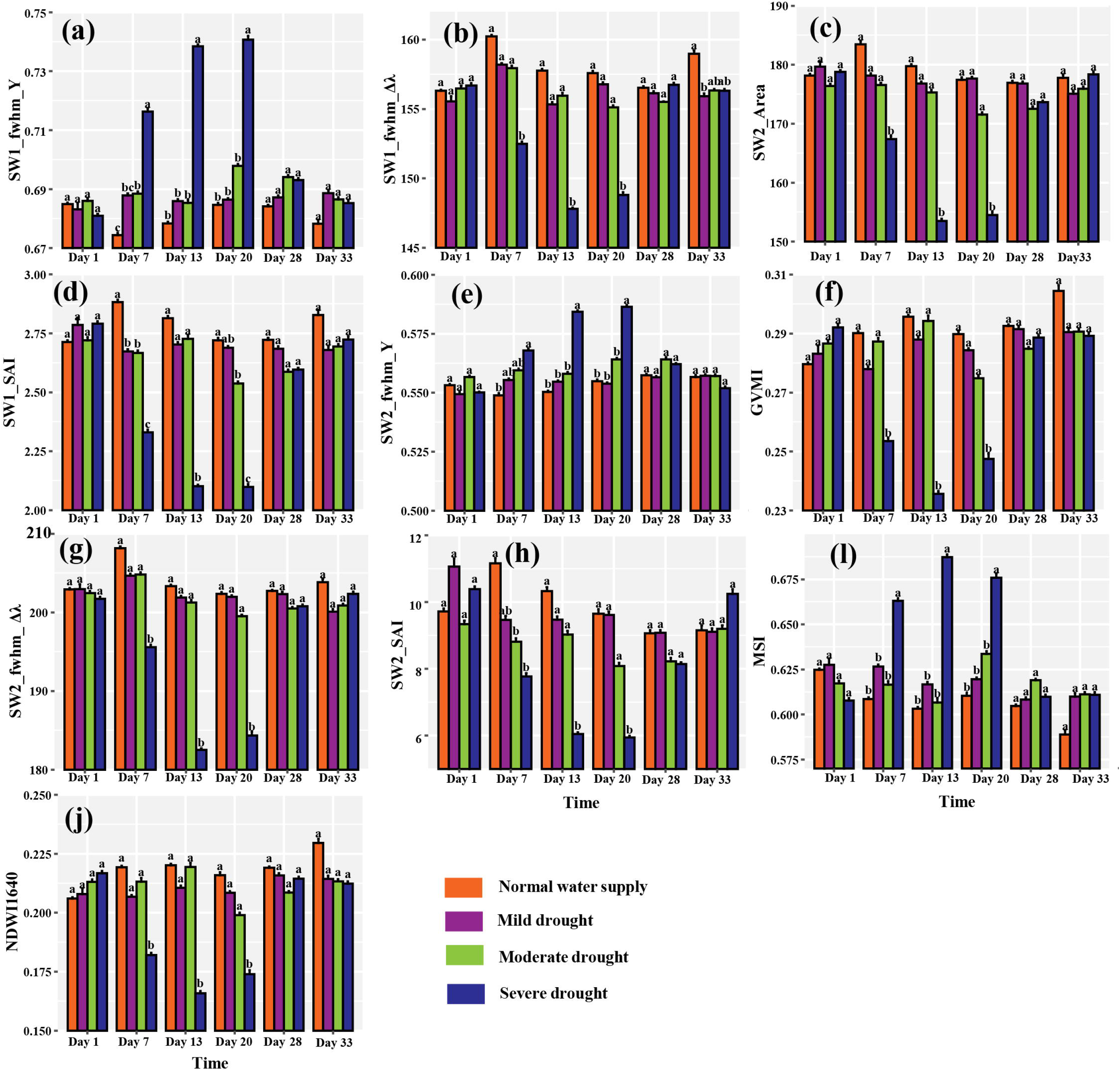
Performance of spectral parameters for revealing the difference of four drought treatments. a, b, c, d, e, g, h, i, j, k and l were indicated as SW1-fwhmY1, SW1-fwhm-Δλ, SW2_area, SW1-SAI, SW2-fwhm-Y, GVMI, SW2-fwhm-Δλ, SW2-SAI, MSI, and NDWI1640.

GI, CI_730_, CI_709_, CIG, NDRE, Rg, CARI, MCTI, PRI, VIS-λ_2_, VIS-λ_*p*_, VIS-Area, VIS-symmetry, VIS-slope, VIS-fwhm X_1_, VIS-fwhm-Δλ, SW1-λ_1_, SW2-λ_1_, SW2-slope, SW2-fwhm-X_1_, C, λ, λ_*p*_, σ and REP of responses to drought were lagging, and spectral variations of different treatments could still be presented in the initial stage of rewatering (i.e. day 28), but great uncertainty occurred (Table 2).

**Table 2:**
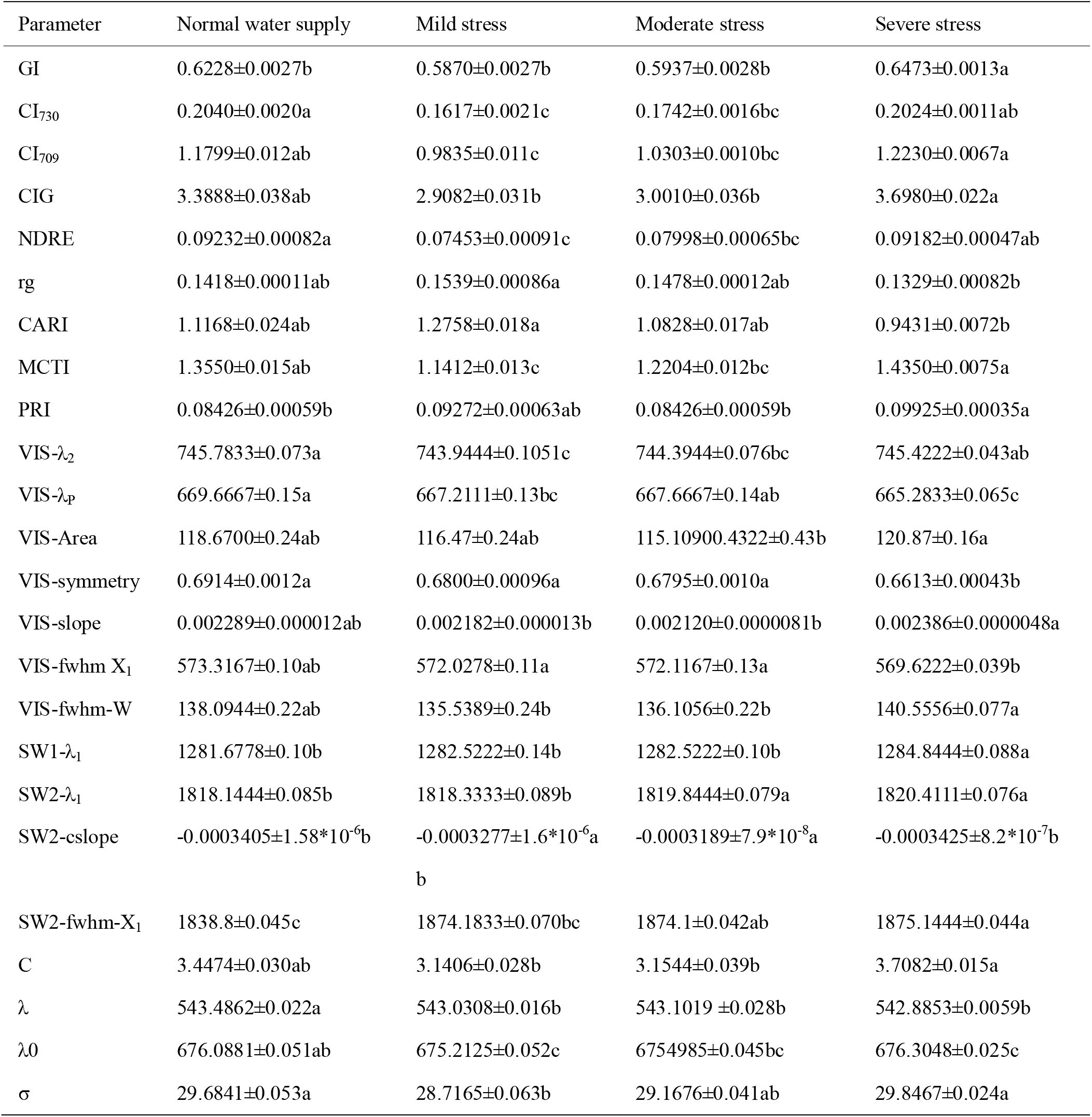
Performance of spectral parameters of normal water supply, mild stress, moderate stress and severe stress in the initial stage of rewatering (i.e., 4-Sep). The data was presented in the form of mean±standard error, significant differences were indicated by different letters in the same row.

### Machine learning algorithms to predict *Pn, Cond* and *Trmmol*

The photosynthetic parameters returned heterogeneous and non-parametric results for the analyzed leaves (Table 3). The analysis showed that *Pn, Cond* and *Trmmol* had high variability and uniform distribution. This heterogeneous dataset was is very useful for building prediction models using machine learning algorithms. Four machine learning algorithms were applied to estimate *Pn, Cond* and *Trmmol* values of lemon leaves. A comparison of the four machine learning algorithms showed that RF demonstrated the best regression performance in terms of *Pn, Cond* and *Trmmol* values. The R^2^ value ranged from 0.88 to 0.92, and the RMSE was 1.86, 0.049 and 1.88 for *Pn, Cond* and *Trmmol*, respectively. The AdaBoost achieved the second highest accuracy except for the *Trmmol*. In the AdaBoost regression models, the R^2^ ranged from 0.49 to 0.69 and the RMSE was 1.84, 0.056 and 2.078 for *Pn, Cond* and *Trmmol*. SVM obtained a moderate performance and presented R^2^ values from 0.28 to 0.64 (Table 4).

**Table 3:** Descriptive data from the photosynthetic parameters’ analysis of the lemon leaves.

| Summary | $Pn$ ($\mu\text{mol CO}_2 \text{ m}^{-2} \text{ s}^{-1}$) | $Cond$ ($\text{mol HO}_2 \text{ m}^{-2} \text{ s}^{-1}$) | $Trmmol$ ($\text{mmol HO}_2 \text{ m}^{-2} \text{ s}^{-1}$) |
| --- | --- | --- | --- |
| Mean | 4.53 | 0.089 | 3.00 |
| Std. Dev. | 2.85 | 0.069 | 2.21 |
| Median | 4.15 | 0.073 | 2.58 |
| Mix. | 12.51 | 0.36 | 11.61 |
| Min. | 0.022 | 0.0014 | 0.037 |
| Coeff. Var. | 63.02 | 77.79 | 73.70 |

**Table 4:** The machine learning algorithms’ accuracy performance for the reflectance data.

|  |  | AdaBoost | GDBoost | RF | SVM |
| --- | --- | --- | --- | --- | --- |
| *Pn* | $R^2$ | 0.69 | 0.99 | 0.92 | 0.64 |
|  | RMSE | 1.84 | 2.55 | 1.86 | 1.91 |
|  | MAE | 1.53 | 1.95 | 1.51 | 1.53 |
| *Cond* | $R^2$ | 0.52 | 0.99 | 0.89 | 0.28 |
|  | RMSE | 0.056 | 0.068 | 0.049 | 0.055 |
|  | MAE | 0.045 | 0.049 | 0.038 | 0.045 |
| *Trmmol* | $R^2$ | 0.49 | 0.99 | 0.88 | 0.50 |
|  | RMSE | 2.078 | 2.48 | 1.88 | 2.06 |
|  | MAE | 1.65 | 1.74 | 1.34 | 1.46 |

To ascertain the relationship between observed and predicted *Pn, Cond* and *Trmmol*, their regression values were plotted (Figure 7). GDboost did not show similarity to a 1:1 relationship (dashed-line—Figure 7b, 7f and 7g). Predictions of Adaboost, RF and SVM were comparatively well related to the observed *Pn, Cond* and *Trmmol* values (Figure 7).

**Figure 7.**
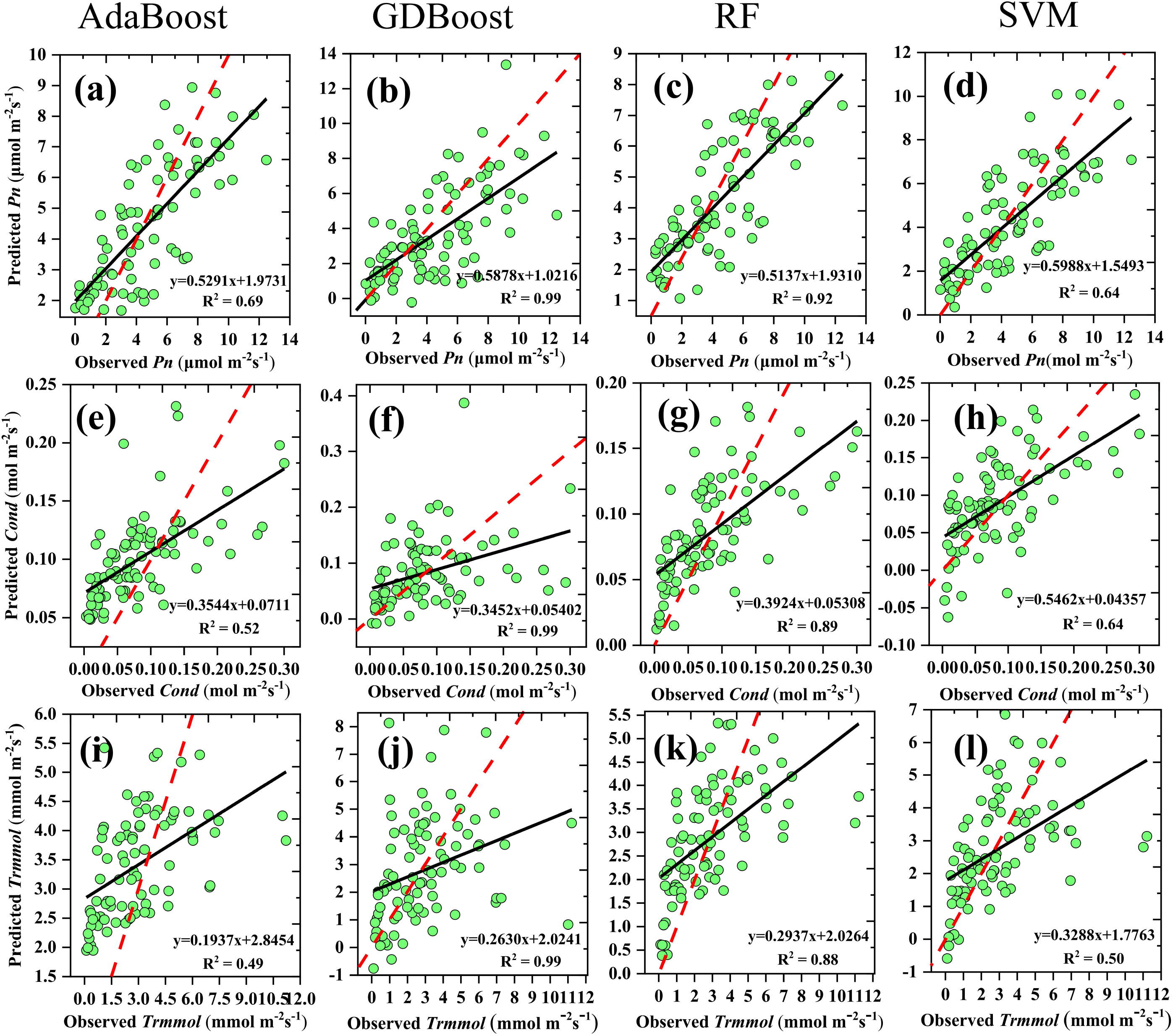
Measured vs. predicted values after applying AdaBoost, GDBoost, random forest (RF) and support vector machine (SVM) model to predict leaf photosynthetic CO_2_ assimilation rate (*Pn* in µmol m^-2^ s^-1^), leaf stomatal conductance (*Cond* in mol m^-2^ s^-1^), and leaf transpiration rate (*Trmmol* in µmol m^-2^ s^-1^). The red line is the 1:1 line and the black line is fitting line between observed and predicted values. (a), (b), (c), (d), (e), (f), (g), (h) and (i) were *Pn*- AdaBoost, *Pn*- GDBoost, *Pn*- RF, *Pn-* SVM, *Cond-* AdaBoost, *Cond-* GDBoost, *Cond-* RF, *Cond-* SVM, *Trmmol*- AdaBoost, *Trmmol*- GDBoost, *Trmmol*- RF and *Trmmol*- SVM.

## Discussion

Global warming leads to increasing drought problems and food shortages. Water deficit monitoring and yield estimation for citrus trees of large area is very critical. Laboratory method of water stress and gas exchange measurements still are cumbersome experimental techniques and not suitable for large-scale monitoring in a short time (Fu et al. 2019, Fu et al. 2020). Novel techniques are thus required to efficiently select for water stress and photosynthetic capacity (Heckmann et al. 2017). Drought resistance is a combination of physiological and biochemical adaptations (Boochs et al. 1990, Ashraf 2010), which can be reflected in the plants’ spectral signature (Zovko et al. 2019). To this end, we studied photosynthetic and spectral responses under water stress, analyzed the properties of leaf reflectance measurements and built corresponding predictive photosynthesis models. It will play an important role in predicting yield and mitigating the impact of drought on yield in a large-scale orchard. To understand the general properties and diversity of leaf reflectance spectra in different drought treatments, we estimated their distribution. Percentage of 97.55 of raw spectra of three water stress was contained in the first three principal components (Figure 2), which revealed a low and unexpected diversity of spectral properties. Five high correlated band regions were identified. This feature of the spectra poses a challenge for the development of robust predictive models. Machine learning algorithms may be urgent to predict photosynthetic characterization because of their advantage of solving multi-collinearity.

Hyperspectral technology can accurately obtain the fine spectral information of plants for accurately monitoring the growth, physiological and biochemical characteristics of plants. Although using remote sensing to assess responses to drought is a very active topic of research, most studies to date have focused on estimating biochemical and structural parameters related to water stress (Sobejano-Paz et al. 2020). In this study, physiological and spectral responses to soil drought were assessed. For citrus, a significant decline of leaf *Pn, Cond* and *Trmmol* was observed after trees suffered water stress (Figure 2, Supplementary Table S1-S2). An effective and alternative method was provided to identify drought stress and its severity early in citrus trees. PRI, NDVI, RVI, GI, C, NMDI, VIS-λ_*p*_, SW1-fwhmX_1_ and SW1-fwhmX_2_ were effective in tracking continuous drought responses in citrus at the beginning of drought treatment when leaves did not show any morphological changes. Once drought stress occurs, leaves quickly close the stomata to reduce water loss, stomatal conductance and transpiration (Sun et al. 2013). Drought stress affects morphological characteristics such as leaf relative water content and leaf area, leaf relative conductivity (Dong et al. 2021). NDVI, RVI, GI, NMDI, SW1-fwhmX_1_ and SW1-fwhmX_2_ have indirect relationships to plant physiological and structural parameters such as water content and greenness (Govender et al. 2009). Especially, PRI was a key remote sensing index, which was surprisingly more sensitive to an early plant water-stress stadium than traditional SVIs from beginning to end and can serve as a pre-visual and continuous water-stress indicator (Suarez et al. 2009, Panigada et al. 2014). This result was contributed the fact that PRI was closely linked to photosynthetic process due to faster changes in xanthophyll pigments comparing other SVIs under stress conditions (Gerhards et al. 2019). Leaf stomatal closing responding to water stress was early than change of leaf morphology and pigment. PRI nearly presented the same respond as photosynthetic parameters with water stress (Figure 2, Table S1, Table S2, and Figure 5a).

In addition to photosynthesis and moisture reduction, the total soluble sugar, soluble protein and starch content increase, whereas chlorophyll a and b content decrease significantly with the extension of the drought period (Dong et al. 2021). SW1-fwhmY, SW1-fwhm-Δλ, SW1-SAI, SW2-fwhm-Y, SW2-fwhm-Δλ, SW2-SAI, SW2-area, MSI, NDWI1640 and GVMI of severe drought were also different significantly than other water stress from the 7th day of drought treatment to the end (Figure 6). SW1 and SW2 was around 1230-1650nm and 1800-2200nm which were highly sensitive to leaf water content. Zovko et al. (2019) also showed SWIR was effective to determine drought stress and its severity in grapevines. Drought led to the change of leaf water, cellulose, starch and lignin content (Zaher-Ara et al. 2016) which was linked to SW1 and SW2(Zhao 2003). Water deficits affect citrus physiology and citrus exposed to drought stress had a higher amount of soluble sugar and a lower amount of starch. The accumulation of soluble sugar and proline indicates a possible role of these osmolytes in drought tolerance (Arbona et al. 2005, Zaher-Ara et al. 2016). MSI, NDWI1640 and GVMI were measured with around 800 and 1600 bands which were just related to starch and sugar. GI, CI_730_, CI_709_, CIG, NDRE, Rg, CARI, MCTI, PRI, VIS-λ_2_, VIS-abs, VIS-Area, VIS-symmetry, VIS-slope, VIS-fwhm X1, VIS-fwhm-Δλ, SW1-λ_1_, SW2-λ_1_, SW2-slope, SW2-fwhm-X_1_, C, λ, λ0, σ, REP of different treatments could still be presented in the initial stage of rewatering. They were related to content of xanthophyll content, chlorophyll, carotenoid, sugar and starch (Zhao 2003, Govender et al. 2009). There was a process of plant recovery, and these indicators may reflect differences in the process of recovery from the drought conditions. It was proved that hyperspectral SVIs and spectral absorption and wavelength position variables was effective in drought stress identification. The limitation was that most of hyperspectral parameters can only distinguish severe drought from all water stress.

To obtain quantitative assessment of water stress and yield prediction, we systematically developed *Pn, Cond* and *Trmmol* prediction models with high precision and evaluated the performance of a variety of models. This was the first time that photosynthetic parameters related to citrus yield had been predicted. In citrus, past studies on predicting physiological traits of citrus mainly focused on nutrient or micronutrient content like N, P, K, Mg, S, Cu, Fe, Mn, and Zn (Osco et al. 2019, Osco et al. 2020). In this study, RF was the best predictor, followed by AdaBoost and SVM (Table 4). RF models had the highest R^2^ (0.92), lower RMSE (1.86) and MAE (1.51). Random forest has been reported to bring high accuracy of physiological traits in crops and forests (Zhao et al. 2019, Osco et al. 2020). Partial least-squares regression model relating to hyperspectral reflectance also has been used to estimate photosynthetic capacity of crops, such as Maize (Yendrek et al. 2017) and tobacco (Meacham-Hensold et al. 2019), however, it did not return higher prediction accuracy than this study. This was contributed to advantage of modeling data in a non-linear and a non-parametric manner and solving the problem of multiple collinearities. Although SVM has advantage of handling high dimensionality data and do well with a limited training dataset(Ma et al. 2019), it performed poorly in comparison with RF in this study. GDboost exhibited severe overfitting and returned unreliable prediction although literature reported GDBoost model performed better RF when estimating forest coverage (Zhou et al. 2020a). It may be concluded that GDBoost present major flaws in modeling highly correlated hyperspectral data.

## Conclusion

Non-destructive and rapid methods for accurate pre-visual water-stress detection and photosynthetic parameters’ estimation was necessary to yield increase and quality improvement of citrus. Photosynthetic parameters presented significant decrease under water stress and this trend was more obviously in upper layer. Original reflectance spectra of three drought treatments presented a low and unexpected diversity. PRI is more sensitive to an early plant water-stress stadium than traditional SVIs and can serve as a pre-visual, persistent and stable water-stress indicator interestingly. Spectral absorption features in SW1 and SW2 regions, MSI, NDWI1640 and GVMI were useful to distinguish severe drought treatment effectively. Photosynthetic rate could be estimated with highest precision by applying hyperspectral leaf reflectance and RF models compared SVM, AdaBoost and GDBoost. To our knowledge, this is one of the first applications of using hyperspectral parameters as indicator for water stress and input for the retrieval of photosynthetic traits in citrus and provides a basis for extending the analysis to other observing platforms, such as unmanned aerial vehicle and satellite data for water condition monitoring and yield increasing quickly and precisely in large-scale orchards.

## Acknowledgments

We would like to thank Prof. Jinzhi Zhang from Huazhong Agricultural University for experiment conduction and data analysis, which greatly improved the manuscript. This research was funded by the national key research and development plan (Grant number, 2019YFD1000104) and project 2662020YLPY020 supported by the fundamental research funds for the central university. This study was also supported by national natural fund project (Grant number, 31901963).

## Author Contributions

J.J. Z. proposed the idea, designed the experiment, organized the writing, and revised the paper. Y.Y.D. supervised the work and analyzed the experimental data. Y.H.Z., Z.M.H., X.Y.Z., and Y.F. J. measured all photosynthetic and spectral data. C.G.H. provided citrus trees as experimental materials and helped to design experiments. All authors have read and agreed to the published version of the manuscript.

## Data availability statement

Data sharing is not applicable to this article as all data is already contained within this article or in the supplementary material.

## Supplementary material

**Supplementary table S1:** One-way ANOVA test results of stomatal conductance (*Cond*, mol m^-2^s^-1^) of upper layer, middle layer, and lower layer in different water stress. The data was presented in the form of mean ± standard error, significant differences were indicated by different letters in the same column.

**Supplementary table S2:** One-way ANOVA test results of leaf transpiration rate (*Trmmol*, mmol m-2s-1) of upper layer, middle layer, and lower layer in different water stress. The data was presented in the form of mean ± standard error, significant differences were indicated by different letters in the same column.

## References

Acevedo E, Hsiao TC, Henderson DW. 1971. Immediate and subsequent growth responses of maize leaves to changes in water status. Plant Physiology 48, 631–636.

Arbona V, Iglesias DJ, Jacas J, Primo-Millo E, Talon M, Gomez-Cadenas A. 2005. Hydrogel substrate amendment alleviates drought effects on young citrus plants. Plant and Soil 270, 73–82.

Ashraf M. 2010. Inducing drought tolerance in plants: Recent advances. Biotechnology Advances 28, 169–183.

Asner GP. 1998. Biophysical and biochemical sources of variability in canopy reflectance. Remote Sensing of Environment 64, 234–253.

Ballester C, Zarco-Tejada PJ, Nicolas E, Alarcon JJ, Fereres E, Intrigliolo DS, Gonzalez-Dugo V. 2018. Evaluating the performance of xanthophyll, chlorophyll and structure-sensitive spectral indices to detect water stress in five fruit tree species. Precision Agriculture 19, 178–193.

Boochs F, Kupfer G, Dockter K, Kuhbauch W. 1990. Shape of the red edge as vitality indicator for plants. International Journal of Remote Sensing 11, 1741–1753.

Boretti A, Rosa L. 2019. Reassessing the projections of the World Water Development Report. Npj Clean Water 2,15. DOI: https://doi.org/10.1038/s41545-019-0039-9

Bruning B, Liu H, Brien C, Berger B, Lewis M, Garnett T. 2019. The Development of Hyperspectral Distribution Maps to Predict the Content and Distribution of Nitrogen and Water in Wheat (Triticum aestivum). Frontiers in Plant Science 10.

Ceccato P, Gobron N, Flasse S, Pinty B, Tarantola SJRsoe. 2002. Designing a spectral index to estimate vegetation water content from remote sensing data: Part 1: Theoretical approach. 82, 188–197.

Chaerle L, Van Der Straeten D. 2000. Imaging techniques and the early detection of plant stress. Trends in Plant Science 5, 495–501.

Chen J, Li F, Wang R, Fan Y, Raza MA, Liu Q, Wang Z, Cheng Y, Wu X, Yang F, Yang W. 2019. Estimation of nitrogen and carbon content from soybean leaf reflectance spectra using wavelet analysis under shade stress. Computers and Electronics in Agriculture 156, 482–489.

Chen X-q, Yang Q, Han J-y, Lin L, Shi L-s. 2020. Estimation of Winter Wheat Leaf Water Content Based on Leaf and Canopy Hyperspectral Data. Spectroscopy and Spectral Analysis 40, 891–897.

Chica EJ, Albrigo LG. 2013. Expression of Flower Promoting Genes in Sweet Orange during Floral Inductive Water Deficits. Journal of the American Society for Horticultural Science 138, 88–94.

Cotrozzi L, Couture JJ, Cavender-Bares J, Kingdon CC, Fallon B, Pilz G, Pellegrini E, Nali C, Townsend PA. 2017. Using foliar spectral properties to assess the effects of drought on plant water potential. Tree Physiology 37, 1582–1591.

Cotrozzi L, Peron R, Tuinstra MR, Mickelbart MV, Couture JJ. 2020. Spectral Phenotyping of Physiological and Anatomical Leaf Traits Related with Maize Water Status. Plant Physiology 184, 1363–1377.

Dash J, Curran P. 2004. The MERIS terrestrial chlorophyll index. International journal of remote sensing, 25, 5403–5413.

Dong T, Xi L, Xiong B, Qiu X, Huang S, Xu W, Wang J, Wang B, Yao Y, Duan C, Tang X, Sun G, Wang X, Deng H, Wang Z. 2021. Drought resistance in Harumi tangor seedlings grafted onto different rootstocks. Functional Plant Biology. DOI: https://doi.org/10.1071/FP20242

Driever SM, Lawson T, Andralojc PJ, Raines CA, Parry MAJ. 2014. Natural variation in photosynthetic capacity, growth, and yield in 64 field-grown wheat genotypes. Journal of Experimental Botany 65, 4959–4973.

Fitzgerald G, Rodriguez D, O’Leary G. 2010. Measuring and predicting canopy nitrogen nutrition in wheat using a spectral index-The canopy chlorophyll content index (CCCI). Field Crops Research 116, 318–324.

Fu P, Meacham-Hensold K, Guan K, Bernacchi CJ. 2019. Hyperspectral Leaf Reflectance as Proxy for Photosynthetic Capacities: An Ensemble Approach Based on Multiple Machine Learning Algorithms. Frontiers in Plant Science 10.

Fu P, Meacham-Hensold K, Guan K, Wu J, Bernacchi C. 2020. Estimating photosynthetic traits from reflectance spectra: A synthesis of spectral indices, numerical inversion, and partial least square regression. Plant Cell and Environment 43, 1241–1258.

Gamon J, Penuelas J, Field CJRSoe. 1992. A narrow-waveband spectral index that tracks diurnal changes in photosynthetic efficiency. Remote Sensing of Environment 41, 35–44.

Gamon JA, Somers B, Malenovsky Z, Middleton EM, Rascher U, Schaepman ME. 2019. Assessing Vegetation Function with Imaging Spectroscopy. Surveys in Geophysics 40, 489–513.

Gao BC. 1996. NDWI - A normalized difference water index for remote sensing of vegetation liquid water from space. Remote Sensing of Environment 58, 257–266.

Garriga M, Romero-Bravo S, Estrada F, Escobar A, Matus IA, del Pozo A, Astudillo CA, Lobos GA. 2017. Assessing Wheat Traits by Spectral Reflectance: Do We Really Need to Focus on Predicted Trait-Values or Directly Identify the Elite Genotypes Group? Frontiers in Plant Science 8.

Gerhards M, Schlerf M, Mallick K, Udelhoven T. 2019. Challenges and Future Perspectives of Multi-/Hyperspectral Thermal Infrared Remote Sensing for Crop Water-Stress Detection: A Review. Remote Sensing 11.

Gitelson AA, Keydan GP, Merzlyak MN. 2006. Three-band model for noninvasive estimation of chlorophyll, carotenoids, and anthocyanin contents in higher plant leaves. Geophysical Research Letters 33.

Govender M, Dye PJ, Weiersbye IM, Witkowski ETF, Ahmed F. 2009. Review of commonly used remote sensing and ground-based technologies to measure plant water stress. Water Sa 35, 741–752.

Heckmann D, Schlueter U, Weber APM. 2017. Machine Learning Techniques for Predicting Crop Photosynthetic Capacity from Leaf Reflectance Spectra. Molecular Plant 10, 878–890.

Huete A, Justice C, Liu HJRSoe. 1994. Development of vegetation and soil indices for MODIS-EOS. 49, 224–234.

Hunt ER, Rock BN. 1989. Detection of changes in leaf water-content using near-infraed and middle-infrared reflectances. Remote Sensing of Environment 30, 43–54.

CFJE Jordan. 1969. Derivation of leafiarea index from quality of light on the forest floor. Ecology 50, 663–666.

Kim MS, Daughtry C, Chappelle E, McMurtrey J, Walthall C. 1994. The use of high spectral resolution bands for estimating absorbed photosynthetically active radiation (A par).

Kong W, Huang W, Zhou X, Mortimer H, Ma L, Tang L, Li C. 2019. Estimating leaf water content at the leaf scale in soybean inoculated with arbuscular mycorrhizal fungi from in situ spectral measurements. International Journal of Agricultural and Biological Engineering 12, 149–155.

Li J-X, Hou X-J, Zhu J, Zhou J-J, Huang H-B, Yue J-Q, Gao J-Y, Du Y-X, Hu C-X, Hu C-G Zhang J-Z. 2017. Identification of Genes Associated with Lemon Floral Transition and Flower Development during Floral Inductive Water Deficits: A Hypothetical Model. Frontiers in Plant Science 8.

Ling S, Dandan S, Yi L, Xiaohua S, Jinqiu W, Tao L, Yunliu Z, Juan X, Xiuxin D, Yunjiang C. 2017. Exogenous gamma-aminobutyric acid treatment affects citrate and amino acid accumulation to improve fruit quality and storage performance of postharvest citrus fruit. Food Chemistry 216, 138–145.

Liu S-R, Zhou J-J, Hu C-G, Wei C-L, Zhang J-Z. 2017. MicroRNA-Mediated Gene Silencing in Plant Defense and Viral Counter-Defense. Frontiers in Microbiology 8, 1801.

Long SP, Bernacchi CJ. 2003. Gas exchange measurements, what can they tell us about the underlying limitations to photosynthesis? Procedures and sources of error. Journal of Experimental Botany 54, 2393–2401.

Long SP, Marshall-Colon A, Zhu X-G. 2015. Meeting the Global Food Demand of the Future by Engineering Crop Photosynthesis and Yield Potential. Cell 161, 56–66.

Ma L, Liu Y, Zhang X, Ye Y, Yin G, Johnson BA. 2019. Deep learning in remote sensing applications: A meta-analysis and review. Isprs Journal of Photogrammetry and Remote Sensing 152, 166–177.

Manevski K, Manakos I, Petropoulos GP, Kalaitzidis C. 2011. Discrimination of common Mediterranean plant species using field spectroradiometry. International Journal of Applied Earth Observation and Geoinformation 13, 922–933.

McDowell N, Pockman WT, Allen CD, Breshears DD, Cobb N, Kolb T, Plaut J, Sperry J, West A, Williams DG, Yepez EA. 2008. Mechanisms of plant survival and mortality during drought: why do some plants survive while others succumb to drought? New Phytologist 178, 719–739.

Meacham-Hensold K, Montes CM, Wu J, Guan K, Fu P, Ainsworth EA, Pederson T, Moore CE, Brown KL, Raines C, Bernacchi CJ. 2019. High-throughput field phenotyping using hyperspectral reflectance and partial least squares regression (PLSR) reveals genetic modifications to photosynthetic capacity. Remote Sensing of Environment 231, 111176.

Mehdaoui R, Anane M. 2020. Exploitation of the red-edge bands of Sentinel 2 to improve the estimation of durum wheat yield in Grombalia region (Northeastern Tunisia). International Journal of Remote Sensing 41, 8984–9006.

Morgan KT, Barkataky S, Kadyampakeni D, Ebel R, Roka F. 2014. Effects of Short-term Drought Stress and Mechanical Harvesting on Sweet Orange Tree Health, Water Uptake, and Yield. Hortscience 49, 835–842.

Mountrakis G, Im J, Ogole C. 2011. Support vector machines in remote sensing: A review. Isprs Journal of Photogrammetry and Remote Sensing 66, 247–259.

Ort DR, Merchant SS, Alric J, Barkan A, Blankenship RE, Bock R, Croce R, Hanson MR, Hibberd JM, Long SP, Moore TA, Moroney J, Niyogi KK, Parry MAJ, Peralta-Yahya PP, Prince RC, Redding KE, Spalding MH, van Wijk KJ, Vermaas WFJ, von Caemmerer S, Weber APM, Yeates TO, Yuan JS, Zhu XG. 2015. Redesigning photosynthesis to sustainably meet global food and bioenergy demand. Proceedings of the National Academy of Sciences of the United States of America 112, 8529–8536.

Osco LP, Marques Ramos AP, Faita Pinheiro MM, Saito Moriya EA, Imai NN, Estrabis N, Ianczyk F, de Araujo FF, Liesenberg V, de Castro Jorge LA, Li J, Ma L, Goncalves WN, Marcato Junior J, Creste JE. 2020. A Machine Learning Framework to Predict Nutrient Content in Valencia-Orange Leaf Hyperspectral Measurements. Remote Sensing 12, 906.

Osco LP, Marques Ramos AP, Pereira DR, Saito Moriya EA, Imai NN, Matsubara ET, Estrabis N, de Souza M, Marcato Junior J, Goncalves WN, Li J, Liesenberg V, Creste JE. 2019. Predicting Canopy Nitrogen Content in Citrus-Trees Using Random Forest Algorithm Associated to Spectral Vegetation Indices from UAV-Imagery. Remote Sensing 11, 2925.

Panigada C, Rossini M, Meroni M, Cilia C, Busettoa L, Amaducci S, Boschetti M, Cogliati S, Picchi V, Pinto F, Marchesi A, Colombo R. 2014. Fluorescence, PRI and canopy temperature for water stress detection in cereal crops. International Journal of Applied Earth Observation and Geoinformation 30, 167–178.

Pedregosa F, Varoquaux G, Gramfort A, Michel V, Thirion B, Grisel O, Blondel M, Prettenhofer P, Weiss R, Dubourg V, Vanderplas J, Passos A, Cournapeau D, Brucher M, Perrot M, Duchesnay E. 2011. Scikit-learn: Machine Learning in Python. Journal of Machine Learning Research 12, 2825–2830.

Penuelas J, Pinol J, Ogaya R, Filella I. 1997. Estimation of plant water concentration by the reflectance water index WI (R900/R970). International Journal of Remote Sensing 18, 2869–2875.

Pinter PJ, Hatfield JL, Schepers JS, Barnes EM, Moran MS, Daughtry CST, Upchurch DR. 2003. Remote sensing for crop management. Photogrammetric Engineering and Remote Sensing 69, 647–664.

Pu R, Ge S, Kelly NM, Gong P. 2003. Spectral absorption features as indicators of water status in coast live oak (Quercus agrifolia) leaves. International Journal of Remote Sensing 24, 1799–1810.

Rodriguez-Galiano V, Sanchez-Castillo M, Chica-Olmo M, Chica-Rivas M. 2015. Machine learning predictive models for mineral prospectivity: An evaluation of neural networks, random forest, regression trees and support vector machines. Ore Geology Reviews 71, 804–818.

Rousel J, Haas R, Schell J, Deering D. 1973. Monitoring vegetation systems in the great plains with ERTS. Proceedings of the Third Earth Resources Technology Satellite—1 Symposium; NASA SP-351, 309–317.

Samaniego L, Thober S, Kumar R, Wanders N, Rakovec O, Pan M, Zink M, Sheffield J, Wood EF, Marx A. 2018. Anthropogenic warming exacerbates European soil moisture droughts. Nature Climate Change 8, 421–426.

Santoso H, Tani H, Wang X, Segah H. 2019. Predicting oil palm leaf nutrient contents in kalimantan, indonesia by measuring reflectance with a spectroradiometer. International Journal of Remote Sensing 40, 7581–7602.

Serbin SP, Singh A, McNeil BE, Kingdon CC, Townsend PA. 2014. Spectroscopic determination of leaf morphological and biochemical traits for northern temperate and boreal tree species. Ecological Applications 24, 1651–1669.

Shah SH, Angel Y, Houborg R, Ali S, McCabe MF. 2019. A Random Forest Machine Learning Approach for the Retrieval of Leaf Chlorophyll Content in Wheat. Remote Sensing 11, 920.

Silva-Perez V, Molero G, Serbin SP, Condon AG, Reynolds MP, Furbank RT, Evans JR. 2018. Hyperspectral reflectance as a tool to measure biochemical and physiological traits in wheat. Journal of Experimental Botany 69, 483–496.

Sobejano-Paz V, Mikkelsen TN, Baum A, Mo X, Liu S, Koppl CJ, Johnson MS, Gulyas L, Garcia M. 2020. Hyperspectral and Thermal Sensing of Stomatal Conductance, Transpiration, and Photosynthesis for Soybean and Maize under Drought. Remote Sensing 12, 3182.

Sonobe R, Hirono Y, Oi A. 2020a. Quantifying chlorophyll-aandbcontent in tea leaves using hyperspectral reflectance and deep learning. Remote Sensing Letters 11, 933–942.

Sonobe R, Yamashita H, Mihara H, Morita A, Ikka T. 2020b. Estimation of Leaf Chlorophyll a, b and Carotenoid Contents and Their Ratios Using Hyperspectral Reflectance. Remote Sensing 12, 3265.

Streher AS, Torres RdS, Cerdeira Morellato LP, Freire Silva TS. 2020. Accuracy and limitations for spectroscopic prediction of leaf traits in seasonally dry tropical environments. Remote Sensing of Environment 244, 111828.

Suarez L, Zarco-Tejada PJ, Berni JAJ, Gonzalez-Dugo V, Fereres E. 2009. Modelling PRI for water stress detection using radiative transfer models. Remote Sensing of Environment 113, 730–744.

Sun XP, Yan HL, Kang XY, Ma FW. 2013. Growth, gas exchange, and water-use efficiency response of two young apple cultivars to drought stress in two scion-one rootstock grafting system. Photosynthetica 51, 404–410.

Trenberth KE, Dai A, van der Schrier G, Jones PD, Barichivich J, Briffa KR, Sheffield J. 2014. Global warming and changes in drought. Nature Climate Change 4, 17–22.

Wang H, Lu K, Pu R. 2016. Mapping Robinia pseudoacacia forest health in the Yellow River delta by using high-resolution IKONOS imagery and object-based image analysis. Journal of Applied Remote Sensing 10, 045022.

Wang X, Xu Y, Zhang S, Cao L, Huang Y, Cheng J, Wu G, Tian S, Chen C, Liu Y, Yu H, Yang X, Lan H, Wang N, Wang L, Xu J, Jiang X, Xie Z, Tan M, Larkin RM, Chen L-L, Ma B-G, Ruan Y, Deng X, Xu Q. 2017. Genomic analyses of primitive, wild and cultivated citrus provide insights into asexual reproduction. Nature Genetics 49, 765–772.

Watt MS, Buddenbaum H, Leonardo EMC, Estarija HJ, Bown HE, Gomez-Gallego M, Hartley RJL, Pearse GD, Massam P, Wright L, Zarco-Tejada PJ. 2020. Monitoring biochemical limitations to photosynthesis in N and P-limited radiata pine using plant functional traits quantified from hyperspectral imagery. Remote Sensing of Environment 248, 112003.

Were K, Bui DT, Dick OB, Singh BR. 2015. A comparative assessment of support vector regression, artificial neural networks, and random forests for predicting and mapping soil organic carbon stocks across an Afromontane landscape. Ecological Indicators 52, 394–403.

Wu H, Fu B, Sun P, Xiao C, Liu J-H. 2016. A NAC Transcription Factor Represses Putrescine Biosynthesis and Affects Drought Tolerance. Plant Physiology 172, 1532–1547.

Xiao M, Li Y, Lu B. 2019. Response of Net Photosynthetic Rate to Environmental Factors under Water Level Regulation in Paddy Field. Polish Journal of Environmental Studies 28, 1433–1442.

Yamashita H, Sonobe R, Hirono Y, Morita A, Ikka T. 2020. Dissection of hyperspectral reflectance to estimate nitrogen and chlorophyll contents in tea leaves based on machine learning algorithms. Scientific Reports 10, 17360.

Yan Z, Teng M, He W, Liu A, Li Y, Wang P. 2019. Impervious surface area is a key predictor for urban plant diversity in a city undergone rapid urbanization. Science of the Total Environment 650, 335–342.

Yendrek CR, Tomaz T, Montes CM, Cao Y, Morse AM, Brown PJ, McIntyre LM, Leakey ADB, Ainsworth EA. 2017. High-Throughput Phenotyping of Maize Leaf Physiological and Biochemical Traits Using Hyperspectral Reflectance. Plant Physiology 173, 614–626.

Yordanov I, Velikova V, Tsonev T. 2003. Plant responses to drought and stress tolerance. Bulgarian Journal of Plant Physiology, 187–206.

Zaher-Ara T, Boroomand N, Sadat-Hosseini M. 2016. Physiological and morphological response to drought stress in seedlings of ten citrus. Trees-Structure and Function 30, 985–993.

Zarco-Tejada PJ, Berjón A, López-Lozano R, Miller JR, Martín P, Cachorro V, González M, De Frutos AJRSoE. 2005. Assessing vineyard condition with hyperspectral indices: Leaf and canopy reflectance simulation in a row-structured discontinuous canopy. Remote Sensing of Environment 99, 271–287.

Zhang C, Denka S, Cooper H, Mishra DR. 2018. Quantification of sawgrass marsh aboveground biomass in the coastal Everglades using object-based ensemble analysis and Landsat data. Remote Sensing of Environment 204, 366–379.

Zhang X-Y, Huang Z, Su X, Siu A, Song Y, Zhang D, Fang Q. 2020. Machine learning models for net photosynthetic rate prediction using poplar leaf phenotype data. Plos One 15, e0228645.

Zhao Q, Yu S, Zhao F, Tian L, Zhao Z. 2019. Comparison of machine learning algorithms for forest parameter estimations and application for forest quality assessments. Forest Ecology and Management 434, 224–234.

Zhao Y. 2003. Principles and Methods of Remote Sensing Application Analysis. Beijing: Science Press.

Zhou J, Dian Y, Wang X, Yao C, Jian Y, Li Y, Han Z. 2020a. Comparison of GF2 and SPOT6 Imagery on Canopy Cover Estimating in Northern Subtropics Forest in China. Forests 11, 407.

Zhou J, Zhou Z, Zhao Q, Han Z, Wang P, Xu J, Dian Y. 2020b. Evaluation of Different Algorithms for Estimating the Growing Stock Volume ofPinus massonianaPlantations Using Spectral and Spatial Information from a SPOT6 Image. Forests 11, 540.

Zhou X, Jun S, Yan T, Bing L, Hang Y, Quansheng C. 2020. Hyperspectral technique combined with deep learning algorithm for detection of compound heavy metals in lettuce. Food Chemistry 321, 126503.

Zhu X-G, Long SP, Ort DR. 2008. What is the maximum efficiency with which photosynthesis can convert solar energy into biomass? Current Opinion in Biotechnology 19, 153–159.

Zovko M, Zibrat U, Knapic M, Kovacic MB, Romic D. 2019. Hyperspectral remote sensing of grapevine drought stress. Precision Agriculture 20, 335–347.

